# A deeper understanding of intestinal organoid metabolism revealed by combining fluorescence lifetime imaging microscopy (FLIM) and extracellular flux analyses

**DOI:** 10.1101/771188

**Authors:** Irina A. Okkelman, Nuno Neto, Dmitri B. Papkovsky, Michael Monaghan, Ruslan I. Dmitriev

## Abstract

Stem cells and the niche in which they reside feature a complex microenvironment with tightly regulated homeostasis, cell-cell interactions and dynamic regulation of metabolism. A significant number of organoid models has been described over the last decade, yet few methodologies can enable single cell level resolution analysis of the stem cell niche metabolic demands, in real-time and without perturbing integrity. Here, we studied the redox metabolism of Lgr5-GFP intestinal organoids by two emerging microscopy approaches based on luminescence lifetime measurement – fluorescence-based FLIM for NAD(P)H, and phosphorescence-based PLIM for real-time oxygenation. We found that exposure of stem (Lgr5-GFP) and differentiated (no GFP) cells to high and low glucose concentrations resulted in measurable shifts in oxygenation and redox status. NAD(P)H-FLIM and O_2_-PLIM both indicated that at high ‘basal’ glucose conditions, Lgr5-GFP cells had lower activity of oxidative phosphorylation when compared with cells lacking Lgr5. However, when exposed to low (0.5 mM) glucose, stem cells utilized oxidative metabolism more dynamically than non-stem cells. The high heterogeneity of complex 3D architecture and energy production pathways of Lgr5-GFP organoids were also confirmed by the extracellular flux (XF) analysis. Our data reveals that combined analysis of NAD(P)H-FLIM and organoid oxygenation by PLIM represents promising approach for studying stem cell niche metabolism in a live readout.

## Introduction

Undiscovered knowledge of mechanisms regulating stem cell niche metabolism during quiescence, proliferation and differentiation is a current limitation to the progression of basic cell biology, tissue engineering and regenerative medicine. Various metabolic and mitochondrial function-affecting factors are increasingly being appreciated as core fundamental markers of pathogenesis, immunological defense mechanisms and differentiation. Specifically, in the realm of stem cell differentiation, understanding such energy production mechanisms could establish a tangible link between stem cell proliferative capacity in relation to ageing and related processes ^1-4^. Recently a complex interplay between metabolism of glycolytic Paneth, stem and other differentiating cell types was revealed^5, 6^: lactate produced by Paneth cells is essential for mitochondrial function of stem cells, while mitochondrial pyruvate transport and active mitochondrial respiration regulate the proliferative activity of stem cells residing within the intestinal crypt. At the level of functional intestine, transcription factor-dependent regulation of mitochondrial respiration is also essential for development of this organ ^7, 8^.

Various biological models are popular in the analysis of stem cell niche metabolism: *in vivo* and model organisms, cultured cells and mini-organ or ‘organoid’ cultures ^9-11^. The intestinal organoid ‘mini-gut’ model combines the advantage of producing the entire diversity of epithelial cells including cell niche micro-environment with relatively quick biomass production and ease of handling. Intestinal organoids recapitulate most functions of the native epithelium and are ‘true organoid’ models, in contrast to other types of aggregated cell models ^12^. Their ‘outside-in’ organization (exterior facing basal membrane with the internal lumen area) enable modeling many aspects of normal intestinal epithelial function in a culture dish or under flow bioreactor conditions. Collectively, organoids are amenable for studies of stem cell niche function by high-throughput next generation sequencing, metabolomics and high content microscopy screening, and by ‘traditional’ methods such as quantitative PCR, western blotting or fluorescence microscopy ^13-15^.

Until recently, only few works dealt with assessment of mitochondrial function of the stem cell niche in intestinal organoids ^5, 6, 16-18^. The standard approach is to analyze the extracellular flux, i.e. ‘XF’ ^19, 20^ which informs an oxygen consumption rate (OCR) coupled with extracellular acidification (ECAR), but since this technology was initially targeted towards high-throughput analysis of 2D homogenous cell populations it cannot assess the complexity and heterogeneity of the multiple cell types present in organoid and 3D cultures. Disruptive methods can be also applied to the analysis of the mitochondrial function of stem cell niche, e.g. combining of FACS of fluorescently labeled stem cells with analysis of cell metabolism ^5, 21^.

Emerging microscopy techniques such as light sheet, phosphorescence and fluorescence lifetime imaging microscopy (FLIM) ^22-24^ allow for non-invasive, cell-specific and direct analysis of metabolism within the organoid models. In particular, FLIM method provides quantitative, multi-parameter and live readout, especially when coupled with microsecond-range emitting phosphors in O_2_-PLIM ^25, 26^ or with endogenous chromophores such as NADH and FAD ^27, 28^. Thus, monitoring molecular oxygen, a direct marker of cell hypoxia, can enable visualization of O_2_ gradients and the heterogeneity of organoid oxygenation in primary murine intestinal organoids ^16, 29^. PLIM also enables an estimation of O_2_ consumption - a primary marker of mitochondrial activity ^30^. Although not previously applied to the analysis of stem cell-derived organoids, two-photon excited NAD(P)H-FLIM allowed for direct assessment of the balance between cell redox and energy production pathways within the intestinal crypt ^31, 32^. FLIM and PLIM, individually and/ or combined, can be used in various multi-parameter assays, such as live and real-time tracing of cell proliferation with oxygenation and measurement of extracellular pH fluxes by the biosensor scaffolds ^25, 29, 33, 34^; these methods are therefore highly attractive for imaging of metabolism in stem cell-based and tissue engineered constructs.

The frequently encountered dilemma in selecting appropriate approaches for the study of organoid metabolism has led to variability in reports regarding metabolic regulation in stem cell function ^5, 6^. In order to address this situation, we utilized a well-established and popular model of Lgr5-GFP intestinal organoids ^35, 36^ for comparing real-time methods assessing mitochondrial function, namely: bulk analysis of OCR on a plate reader, PLIM measurement of cellular oxygenation, and two-photon NAD(P)H-FLIM imaging. Our findings demonstrate that O_2_-PLIM and NAD(P)H-FLIM are highly complementary of each other and that, when combined with live cell tracing of stem cells, are powerful methods for the functional studies of stem cell niche metabolism and its cell-cell heterogeneity.

## Results

### Method 1: Analysis of oxygenation by phosphorescence lifetime imaging microscopy (PLIM)

Recently, we have optimized and reported phosphorescence quenching microscopy (O_2_-PLIM) for the profiling of intestinal organoid oxygenation. We demonstrated a small molecule O_2_ probe, Pt-Glc, to be compatible with one-photon PLIM excitation, capable of selectively staining epithelial cells and demonstrating reliable calibration within various biological models ^17, 37-39^. The principle of this method is outlined in Fig. 1A whereby the phosphorescence of Pt-Glc is specifically quenched by molecular oxygen (O_2_) and the decay curves collected via various PLIM imaging techniques (e.g. time-domain, frequency modulation or other means ^40-42^) are used for calculating phosphorescence lifetimes and subsequent calibrations. PLIM enables improved multiplexed tracing of cell types present in the organoid (crypt and villi regions), e.g. by sequential measurement of Lgr5-GFP fluorescence (stem cells) and with other methods, such as labeling proliferating cells by FLIM ^17^. Depending on the structure of O_2_ probe and available instrumentation, PLIM can be performed under conventional laser scanning (one-photon) and two-photon excitation modes ^26^. Fig. 2A shows typical example of the phosphorescence decay for O_2_-sensitive Pt-Glc probe, which can be fit with mono-exponential function.

**Figure 1.**
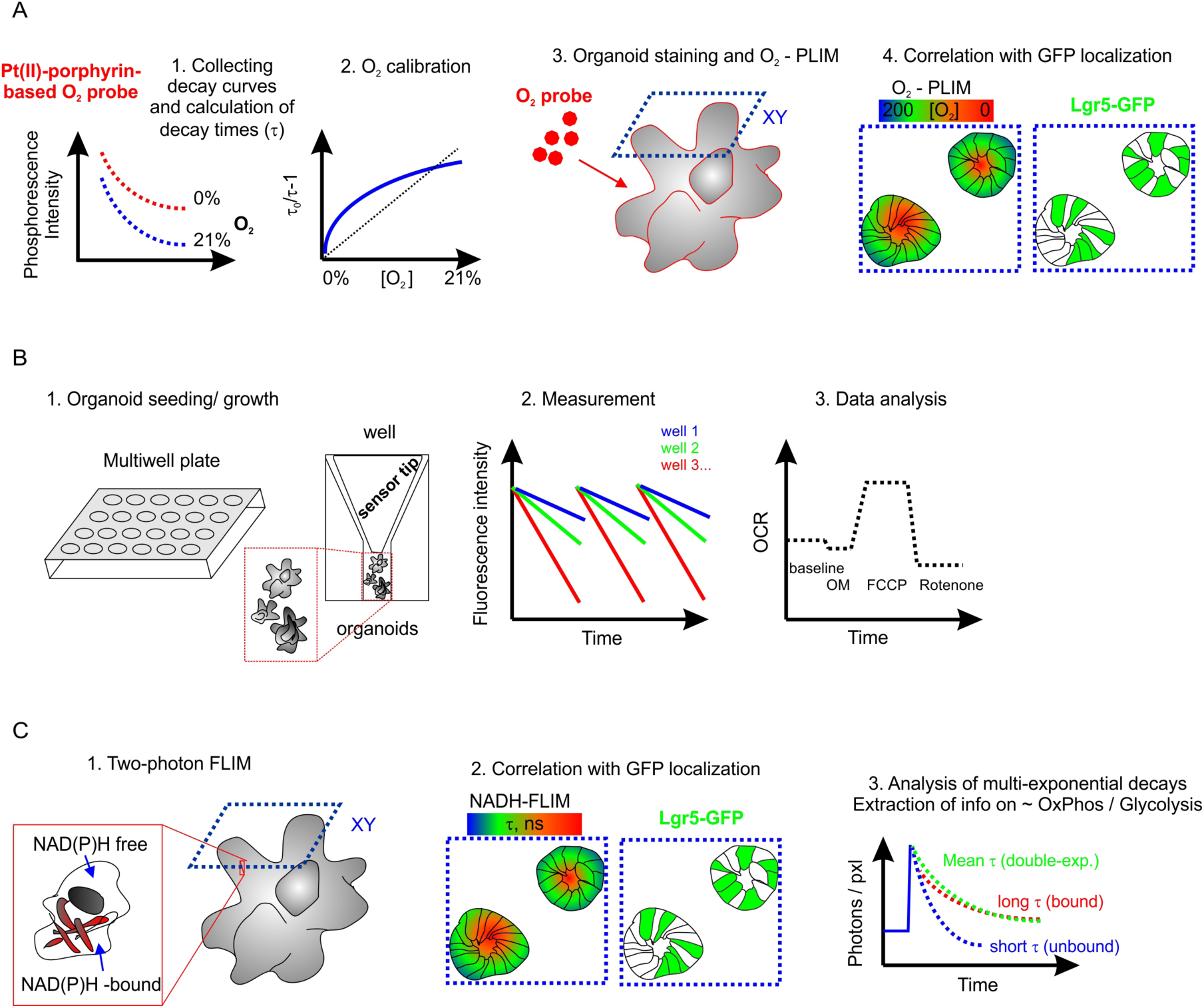
Schematic representation of methods of analysis of the Lgr5-GFP intestinal organoids. A: O_2_-PLIM requires a prior staining of samples with O_2_ probe and specialized PLIM microscope; typically capable of measuring phosphorescence lifetimes in the range of 10-60 μs range of phosphorescence lifetimes. B: Microplate-based extracellular flux (XF) does not need preliminary staining, is well optimized and automated but needs prior seeding of organoids into the assay plates. This method enables only bulk analysis of cell populations. C: NAD(P)H-FLIM relies on the sensitive detection of endogenous NAD(P)H, FAD and other endogenous compounds but needs specialized two-photon FLIM microscope and sophisticated analysis of multi-exponential decay times.

**Figure 2.**
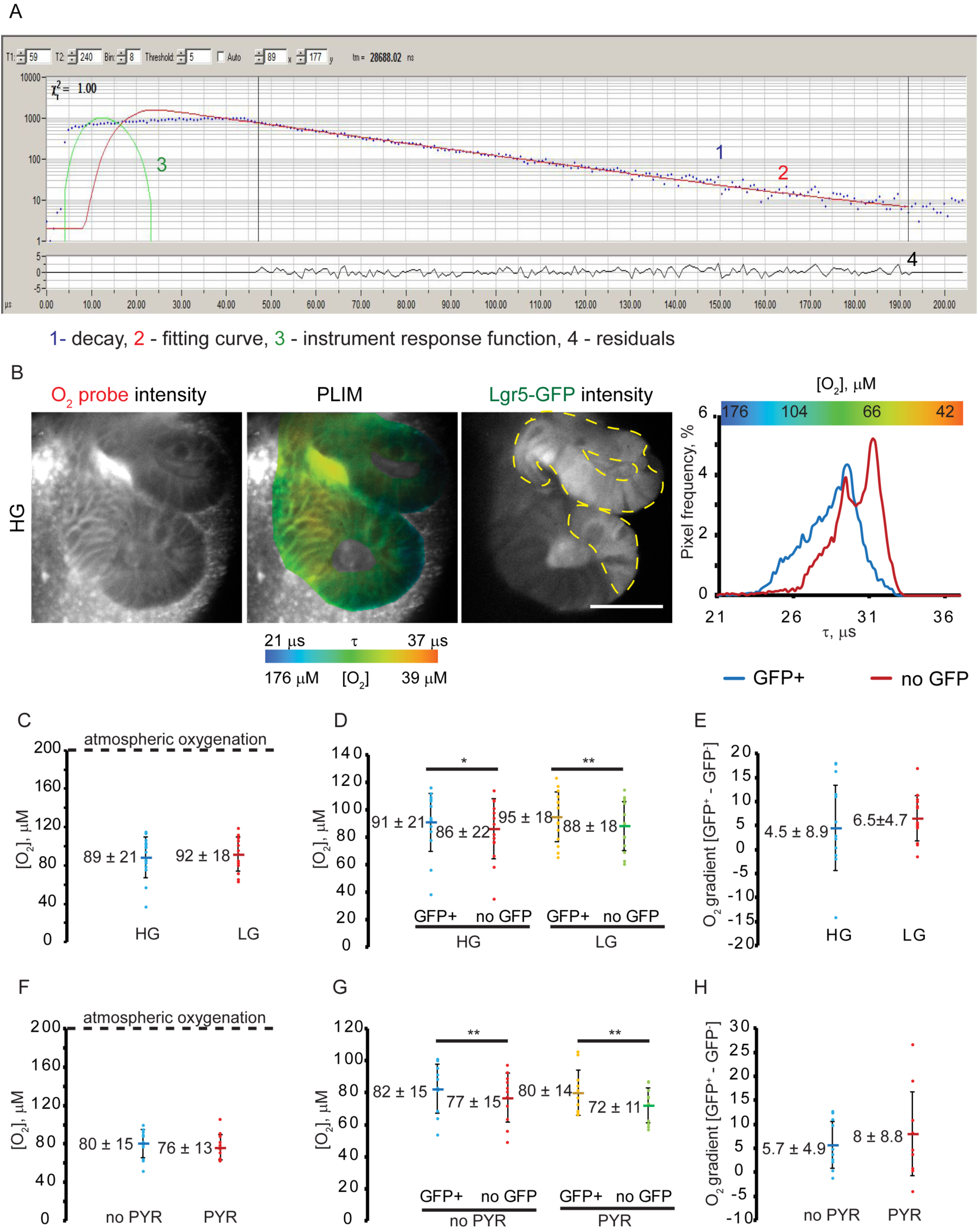
Analysis of Lgr5-GFP organoid oxygenation by PLIM method. A: Typical phosphorescence decay of O_2_ probe in organoids: photons, fitting function, IRF and residuals are indicated by numbers. B: Example of O_2_-PLIM and GFP fluorescence images (single optical sections) obtained in HG media. Scale bar is 50 μm. Phosphorescence lifetime distribution histograms for GFP^+^ and no GFP regions are shown on the right. C-H: O_2_-PLIM analysis of organoids pre-treated in HG (also ‘PYR’ medium), LG and ‘no PYR’ media (1.5 h). Each point indicates average values calculated from individual optical sections. Line marker indicates mean values for each experimental group. Asterisks indicate significant differences (* p<0.1, ** p<0.05). Bars indicate standard deviations. C, F: Effect of media composition on oxygenation of organoids. No significant differences were observed using *t*-test (p=0.1). D, G: Effect of media composition on oxygenation in GFP^+^ and no GFP areas. Differences revealed by using paired *t*-test are indicated (* p<0.1, ** p<0.05). E, H: Effect of media composition on difference in oxygenation between GFP^+^ and no GFP. No significant differences were detected using independent *t*-test (p=0.1).

Within this study we adopted a transgenic Lgr5-GFP-expressing organoid cell line, which enables live tracing of stem cell niche from multipotency to terminal differentiation ^35^. By using previously established methodology ^43^, we marked GFP fluorescence as identifying stem cell niche regions and conversely, regions with low fluorescence as differentiated cells, to perform cell-population specific analysis of O_2_ by PLIM (Fig. 2B). By keeping organoids in ENRVC medium (see Methods section) we maintained them in highly proliferative state, characterized by high number of GFP^+^cells, while switching to EN medium resulted in differentiation/ maturation and a decrease of GFP^+^cell number ^5, 36^. Our preliminary experiments in ENRVC, ENR and EN media found that gradual decrease and loss of GFP expression in ENR- and EN-grown organoids complicates the comparison of GFP^+^and no GFP cells (not shown); we thus focused our study on ENRVC organoids that provided detectable levels of GFP. Organoid cultures were challenged with growth medium containing two different glucose concentrations for 1.5 hours: 0.5 mM (Low Glucose, LG) and 10 mM (High Glucose, HG). Phosphorescent signals from the Matrigel^®^ and luminal regions were not analyzed as the calibration of Pt-Glc within these areas can differ ^17^.

We looked if organoids could rapidly adjust their energy production pathways (glycolysis and oxidative phosphorylation), in response to glucose availability, with oxygenation as a readout parameter. Two independent experiments (see Supplementary Table 1 for individual and combined data) showed a similar trend despite an intrinsically high organoid heterogeneity^16^. For clarity, Fig. 2 presents the data of one experimental replicate. We analyzed global organoid oxygenation by calculating average values for both GFP^+^and no GFP regions in individual optical sections. At basal conditions, organoids displayed moderate deoxygenation of 89-92 μM, corresponding to ∼ 8-9% O_2_ (Fig. 2C), showing no statistical difference between HG and LG groups (p>0.1). Furthermore, we compared the average oxygenation in stem cell niche (calculated from GFP^+^) and neighboring (no GFP) regions in both experimental groups.

Paired *t*-test revealed statistical differences between the GFP^+^and no GFP regions (Fig. 2D) in HG (91 μM O_2_ for GFP^+^ region and 86 μM O_2_ for no-GFP region, p<0.1) as well as in LG-treated organoid populations (95 μM O_2_ for GFP^+^ regions and 88 μM O_2_ for no GFP regions, p<0.05). The comparison of GFP^+^or no GFP areas between HG and LG treatments showed no statistical difference in oxygenation, in response to decreasing glucose content in the media (p>0.1, Fig. 2E). However, in LG medium such difference was increased, potentially due to activation of cell respiration with changing metabolic needs in populations of stem and differentiated cells, being ∼4.5 μM O_2_ for HG vs. 6.5 μM O_2_ for LG (Fig. 2E) and this tendency continued in separate experiments (Table S1) with decreasing p-values with increasing experimental N-number.

Pyruvate metabolism has been identified recently as an important regulator of stem cell function ^6^ and we thus looked at the effect of short-term pyruvate withdrawal from the culture medium containing the high concentration of glucose (Fig. 2F-H). Following the same experimental design and analysis we compared total organoid oxygenation differences between GFP^+^and no GFP areas in absence of (no PYR) and in the presence of sodium pyruvate (PYR). Again, the bulk analysis of the average global oxygenation of organoids did not reveal significant differences (p>0.1) between no-PYR and PYR treatments producing mean values of 80 μM O_2_ and 76 μM O_2_ respectively (Fig. 2F). In addition, the paired *t*-test showed significant differences between GFP^+^and no-GFP cell oxygenation in both no-PYR (82 μM O_2_ vs. 77 μM O_2_, p<0.05) and PYR (80 μM O_2_ vs. 72 μM O_2_, p<0.05) (Fig. 2G). The differences between oxygenation of GFP^+^ vs. no GFP regions showed no statistical decrease in ‘no-PYR’ sample (5.7 μM O_2_ vs. 8 μM O_2_ for no-PYR and PYR groups, p>0.1), potentially indicating an overall decrease of cellular respiration under these conditions (Fig. 2H). However, analysis of combined data from two independent experiments showed no changes in p-value (Table S1) in agreement with hypothesis that the effect of pyruvate withdrawal could be less impactful than glucose deficiency. Indeed, a lack of free pyruvate supplementation could be masked by pyruvate production through glycolysis and glutaminolysis ^28^. In summary, O_2_-PLIM assessment enabled a sensitive detection to discern oxygenation differences between stem cell niche and neighbor mature cells.

### Method 2: Oxygen consumption rate measured by the extracellular flux (XF) assay

Extracellular flux (XF) is today a routinely employed method for assessing oxygen consumption rate (OCR) and additional parameters to enable multi-factorial analyses of mitochondrial function and cell energy budget ^30^. The highly sensitive measurement of OCR at rest and upon pharmacological treatment with metabolic interrogators in a small measurement volume format is possible through the use of built-in solid-state sensors (Fig. 1B). However, the assessment of organoids using XF demands a large amount of samples or crypts, being approx. > 200-300 crypts per experimental point^19^. Therefore, to compare it with O_2_-PLIM which provides analysis of single crypt regions at single cell / organoid resolution, the highly sensitive Agilent^®^ XF96 platform was selected as a comparative tool. Standard flat-bottom XF96 plates were found to be convenient for organoid culture for 3-7 days although standardizing the quantity of plated organoids per well proved difficult, with the best dilution practice achieving between 4 to 21 organoids per well. However, these numbers can be counted and further normalized per GFP fluorescence / cell number or by other means. Figure 3 shows a typical bioenergetic profile for organoids observed on XF96 for ENRVC organoids exposed to low and high glucose conditions, normalized per organoid number per well. This method confirmed strong variability of oxidative metabolism in organoids ^16^ and their low sensitivity to inhibition with oligomycin and rotenone. Still we were able to detect a response to mitochondrial uncoupling by FCCP. This data also demonstrated that XF96 platform had potential to detect flux (OCR) from single organoids.

**Figure 3.**
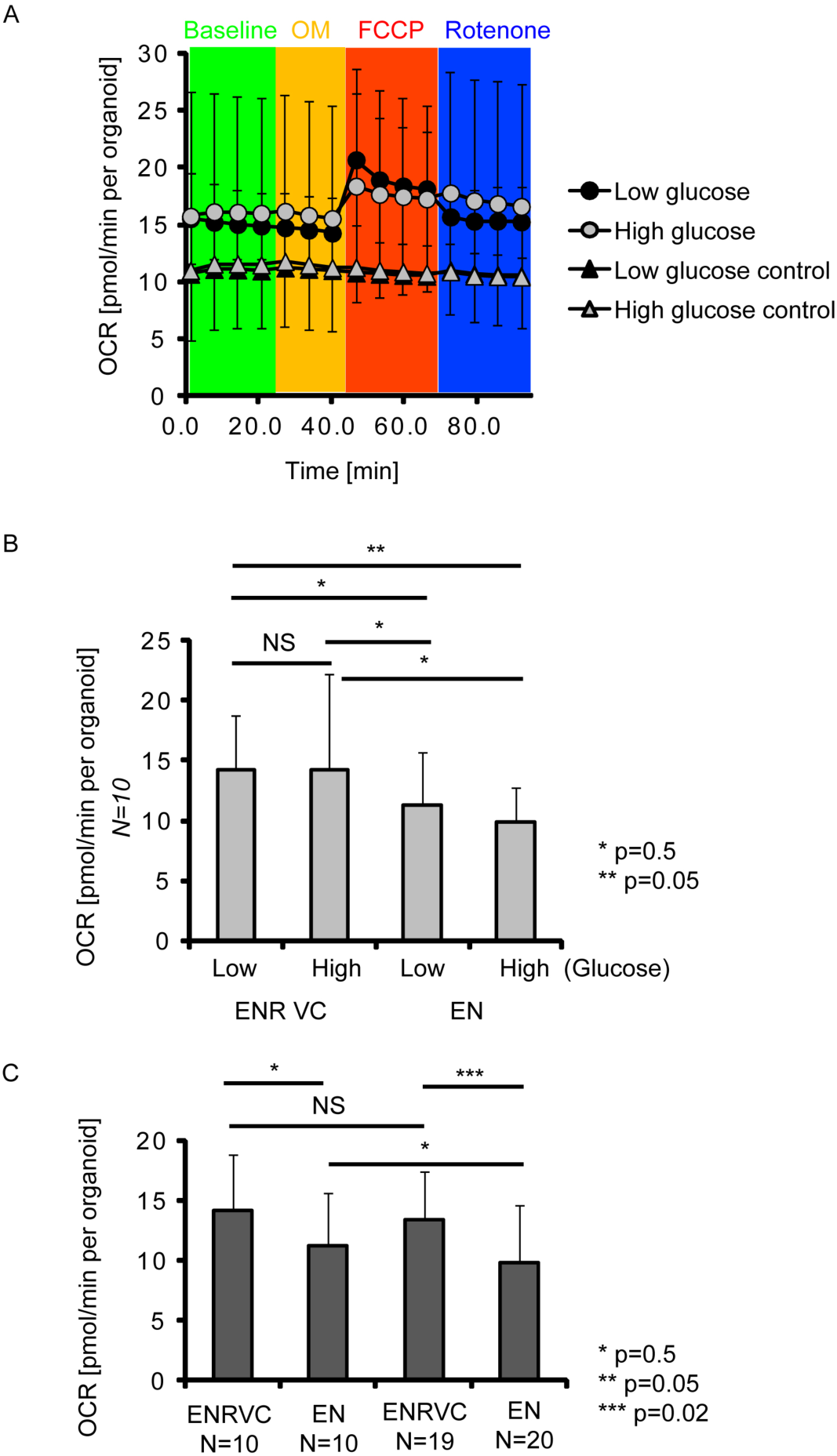
Measurement of Lgr5-GFP organoids oxygen consumption rate by XF96 platform. A: Typical example of OCR profile measured with organoids pre-treated in LG or HG media, with (sequential addition of 1 μM oligomycin, FCCP and rotenone) and without (‘control’) stimulations. Averaged values with standard deviations are shown (N= 10). B: Averaged resting OCR values for organoids grown either in ENRVC or EN media and preincubated in LG or HG media. N=10. C: Effect of increase of N number on the observed differences in OCR. NS-not significant. P values indicate results of independent *t*-test.

As the XF96 platform could provide high throughput and the capability to work with multiple samples, we analyzed the effect of ENRVC and EN media on organoid oxidative metabolism. Interestingly, even with a high variability, we saw overall decrease in OCR upon differentiation conditions (ENR VC vs. EN media). In EN at low glucose conditions organoids also showed slightly increased OCR, potentially due to activation of mitochondria. However, one must keep in mind these data were not normalized to the number of GFP^+^regions or number of crypts. Further, we found that method performance could be improved by increasing a number of replicates (Fig. 3C) or by incorporating more sophisticated means of normalizing per crypt / GFP amounts or by FACS of each measured well after the analysis.

### Method 3: Two-photon excited fluorescence lifetime imaging (FLIM) of NAD(P)H

Autofluorescence originating from excited NAD(P)H can serve as useful redox marker which can be measured non-invasively using two-photon excited FLIM. This enables discrimination between OxPhos and glycolysis in cells ^28, 32, 33^. The principle of this methodology is briefly outlined in Fig. 1C: cells exhibit mixed pools of free (short emission lifetimes, ∼ 400 ps) and enzyme-bound (> 1.5 ns emission lifetimes) NADH and NADPH produced and involved in glycolysis, pentose phosphate pathway, Krebs cycle, oxidative phosphorylation, cytochrome p450 activity and other processes. Spectrally and in the fluorescence lifetime domain it is difficult to separate NADH from NADPH and thus they are often designated as combined pools. Frequently, this method is complemented by measuring FAD (Krebs cycle input) or other compounds ^44^, but its performance depends on the cell type and sensitivity of the available equipment. Here, we focused on simplified two-photon excited NAD(P)H-FLIM imaging as the most frequently used approach in literature.

By using organoids exposed to HG (PYR), LG and no PYR conditions, we performed NAD(P)H-FLIM imaging of GFP^+^and no GFP regions (Fig. 4, Table S2). Noteworthy, intestinal organoids also display profound red autofluorescence of the lumen region, partially originating from products of degradation of cytochrome P450, cell debris and other factors and which can seriously interfere with many fluorescent tracers in ‘intraluminal’ measurements, e.g. glucose uptake tracers ^17, 29^. As a quality control measure, we also measured fluorescence lifetime of GFP and considered regions with τ (GFP) = 2.4±0.1 ns as indicative for its positive expression. NAD(P)H possesses a multi-exponential fluorescence decay, which is normally fitted using a double-exponential curve with mean fluorescence lifetime (τ_m_) derived from two different lifetimes and corresponding fractions (τ_1-2_, α_1-2_). The typical example of NAD(P)H fluorescence decay, its fitting and residuals are shown on Fig. 4A. The short fluorescence lifetime τ_1_ and fraction α_1_ relate to free NAD(P)H, while long fluorescence lifetime τ_2_ and fraction α_2_ correspond to protein-bound NAD(P)H. As τ_1_ values can be affected by the instrument response function, it is difficult to use them for statistical analysis and interpretation of results ^28^. Thus, we focused on analysis of τ_2_ and α_2_ upon different treatment conditions. The results on corresponding τ_m_ are shown on Fig. S3B.

**Figure 4.**
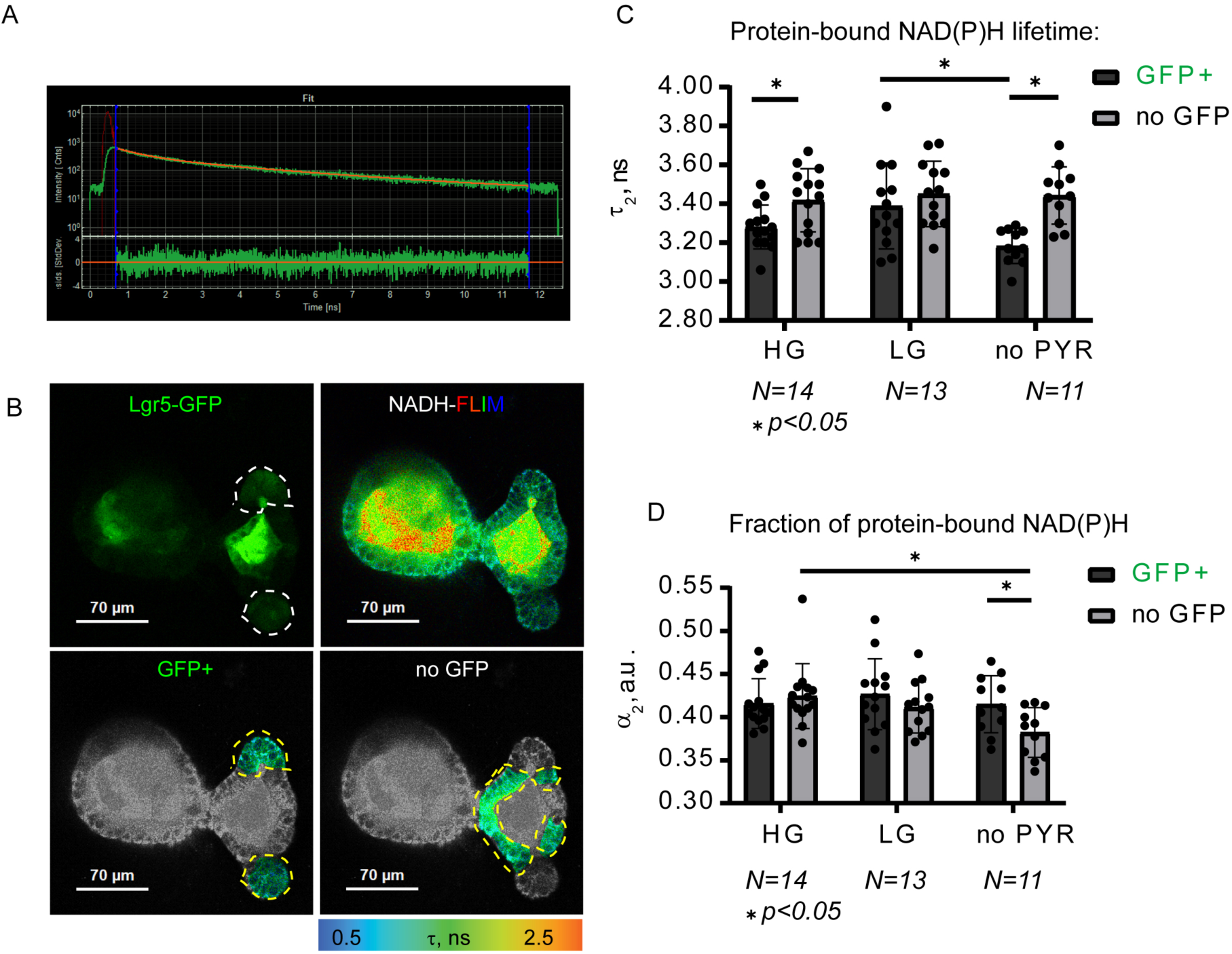
NAD(P)H-FLIM imaging of metabolism in Lgr5-GFP organoids. A: example of multi-exponential decay observed for NAD(P)H signals. B: examples of images of Lgr5-GFP (fluorescence intensity) and NAD(P)H-FLIM for whole organoid and selected GFP^+^ or no GFP regions. C, D: summary of protein-bound lifetimes and the fractions of bound NAD(P)H. Significant differences (paired *t*-test, p<0.05) are indicated with asterisks.

Within HG-treated organoids we detected a significant increase in protein-bound NAD(P)H lifetime (τ_2_) (Fig. 4C) in no GFP regions however the fraction of bound NAD(P)H (α_2_) (Fig. 4D) was not statistically different between the two cell types (GFP^+^ and no GFP) in HG-treated organoids. The average fluorescence lifetime (τ_m_) in differentiated cells exposed to HG showed a significant increase indicating a tendency towards increased oxidative phosphorylation when compared to stem cells (Fig. S3B). However, in LG-treated organoids there were no significant differences in both τ_2_ and τ_m_ (GFP^+^ vs. no GFP cells), suggesting that stem cells could increase their activity of oxidative phosphorylation, while differentiated no GFP cells retained the same activity of oxidative phosphorylation.

When organoids were exposed to ‘no PYR’ conditions, GFP^+^cells again showed both significantly reduced fluorescence lifetimes of protein-bound NAD(P)H (τ_2_) and average fluorescence lifetime (τ_m_) when compared to differentiated no GFP cells. (Fig. S3B). Additionally, differentiated cells had a significantly lower fraction of bound NAD(P)H (α_2_) in contrast with stem cells in no PYR media. Hence, the average lifetime calculated is highly dependent on the fluorescence lifetime of protein bound NAD(P)H. These results potentially indicate a decrease of oxidative phosphorylation activity under no PYR conditions, especially within the differentiated cells.

According to Blacker et al ^28^, a shift of NAD(P)H fluorescence lifetime decays and corresponding τ_2_ (equal to τ_bound_) values does not simply show the alteration of OxPhos / Glycolysis ratio, but primarily reflects a readjustment of the protein bound [NADPH]/[NADH] ratio in cells, where protein-bound NADPH possesses a higher fluorescence lifetime than protein-bound NADH. With this assumption, in comparison to no GFP, GFP^+^ regions had statistically lower τ_2_ (and [NADPH]/[NADH] ratio), which was highly sensitive to glucose and availability of pyruvate (τ_2_ values were 3.28 ns, 3.39 ns and 3.18 ns in HG, LG and no PYR media correspondingly, Fig. 4C). The detailed statistical analysis by paired *t*-test of τ_2_ values between GFP^+^ and no GFP regions at each treatment showed that differences in their [NADPH]/[NADH] ratio increased under no-PYR conditions and decreased to non-significant levels in LG medium (Fig. 4C). Thus, changes in τ_2_ values reflecting [NADPH]/[NADH] ratio for GFP^+^were evident between LG and no-PYR treatment group (p<0.05, Fig. 4C).

In contrast to GFP^+^, no GFP regions had mostly unaffected τ_2_ values by mild metabolic stress (i.e. decrease of glucose, pyruvate withdrawal; τ_2_ values were 3.42 ns, 3.45 ns and 3.44 ns in HG, LG and no PYR media correspondingly, no significant difference at p >0.05) implying that these differentiated and mature intestinal cells tended to keep their [NADPH]/[NADH] ratio at constant levels (Fig. 4C). The α_2_ (equal to α_bound_) parameter informs on NAD(P)H-bound fraction and correlates with the overall proportion of bound NAD(P)H, showing alterations of the protein-bound NAD(P)H pool which is primarily due to shifts in the presence of the free NAD(P)H pool. Thus, we compared α_2_ between GFP^+^ and no GFP regions under HG, LG and no PYR treatments. As anticipated, changes in α_2_ were similar to τ_2_ (α_2_ values were 0.42, 0.43, 0.41 and 0.43, 0.41, 0.38, corresponding to GFP^+^and no GFP regions in HG, LG and no-PYR media), but they were less affected by changes in glucose and pyruvate content, with no significant differences for GFP^+^regions (Fig. 4D, Table S2). However, no GFP regions displayed significant decrease of α_2_ value (p<0.05) in no PYR conditions (Fig. 4D), demonstrating that even if [NADPH]/[NADH] ratio was unchanged, the total pool of reduced protein-bound NAD(P)H decreased. Interestingly, our analysis also revealed no statistical difference for α_2_ between GFP^+^ and no GFP regions (except for no PYR conditions) meaning that generally all cells in organoids had very similar levels of protein-bound NAD(P)H. Thus, NAD(P)H-FLIM analysis showed that in response to changing glucose and pyruvate content, stem cell niche (GFP^+^) and differentiated (no GFP) epithelial cells adapted their metabolism by different means.

## Discussion

### Overall comparison of methodologies

The comparisons described in this study demonstrate the advantages and limitations of each methodology in assessing intestinal organoid metabolism. Extracellular flux (XF) analysis cannot provide single-cell analysis unless the sample is physically detached or dissociated from its substrate, i.e. by enzymatic treatment. On the other hand, XF96 platform was sensitive enough to confirm previously reported heterogeneity of organoid oxygenation ^16^. Indeed, extracellular flux analysis requires improvement for the normalization of heterogeneous cell types and cell mass / densities present in organoids. Microscopy-based FLIM and PLIM methods, while rarely reported together, provided direct readouts of cell metabolism both within and outside Lgr5-GFP-labeled stem cell niches. Potentially, NAD(P)H-FLIM and O_2_-PLIM methods could be multiplexed with each other, and with additional measurement assays, such as labeling of other cell types (genetically or via fluorescent tracers), labeling cell proliferation ^29^ and measurement of mitochondrial membrane potential^18^. Curiously, the majority of commercially available FLIM-PLIM microscopes have limited microsecond PLIM measurement range ^33^, many high-performance O_2_ probes are not suitable for two-photon excitation ^37^, cannot stain organoids or possess other drawbacks ^45, 46^.

An important feature of the microscopy approaches presented here is that they can be performed with live samples and indeed are compatible with downstream assays, such as tissue clearing with light sheet microscopy, coherent anti-Stokes Raman spectroscopy and others ^15, 22^. The aim of our study was not in presenting the most comprehensive or striking measurement example but to show in a simple and easy to reproduce form the examples of FLIM and PLIM uses. Since NAD(P)H-FLIM and O_2_-PLIM separately provided rather limited information, it is clear that they should be used together, when possible.

### Identified methodological limitations of multi-parameter FLIM-PLIM of organoids

A key criteria for success in using FLIM and PLIM lies in a straightforward and clear data processing workflow that can be automated, and largely depends on the fitting of luminescence decays of the intended fluorophores (e.g. NAD(P)H) or phosphors. Without considering potential problems of probe delivery and calibration performance ^45^, most tested phosphorescent probes provide mono-exponential decay fits, well-documented physical interactions with O_2_, ensuring a high precision of measurements and data interpretation ^42, 47^. In contrast, most fluorescent dyes, probes and fluorescent protein biosensors display double or multi-exponential decay fits, which are meaningful but require complex interpretation or the use of phasor approach plotting ^27, 32, 48^. Thus, detailed interpretation of NAD(P)H imaging is not straightforward, often demanding additional pharmacological tests, segmentation based on signal localization and potentially use of additional tracers ^27, 28, 32, 44^.

The progress in automation and improved ability to produce 3D reconstruction of PLIM and NAD(P)H-FLIM images is very important for studies of intestinal organoids and related 3D stem cell-based organoids. The ability to produce fully functional 3D FLIM / PLIM data reconstructions with an option to optically section/ segment in any certain direction (not only in XY but also Z) and subsequently analyze the pixels present in this cross-section is highly important for future studies. Indeed, organoids are multicellular structures of various sizes and shapes and selection of appropriate cross-section for analysis is essential. For example, we observed more striking differences in oxygenation with more ‘typical’ cross sections (Fig. 5A) than with ones having mosaic-like distribution of GFP^+^cells (Fig. 5B). The comparison of GFP^+^and no GFP regions in individual organoids revealed two main types of optical sections: the group including highly significant (independent *t*-test, p<0.05) and nearly significant (independent *t*-test, p<0.1) statistical differences in oxygenation between GFP^+^and no GFP and the group with no statistical differences (p>0.1). This difference was clearly observed on phosphorescence lifetime images and respective O_2_ distribution histograms (Fig. 2B, Fig. S1) and confirmed by Pearson correlation analysis of ROI oxygenation with GFP intensity values (Fig. 5). Even though the number of sections with statistical differences was lower (∼ 41%) than the number of sections without statistically confirmed difference, the calculated oxygenation difference between GFP^+^and no GFP areas in most organoids was positive or slightly negative. These data also indicate GFP^+^regions generally have higher cell oxygenation than no GFP regions in intestinal organoids.

**Figure 5.**
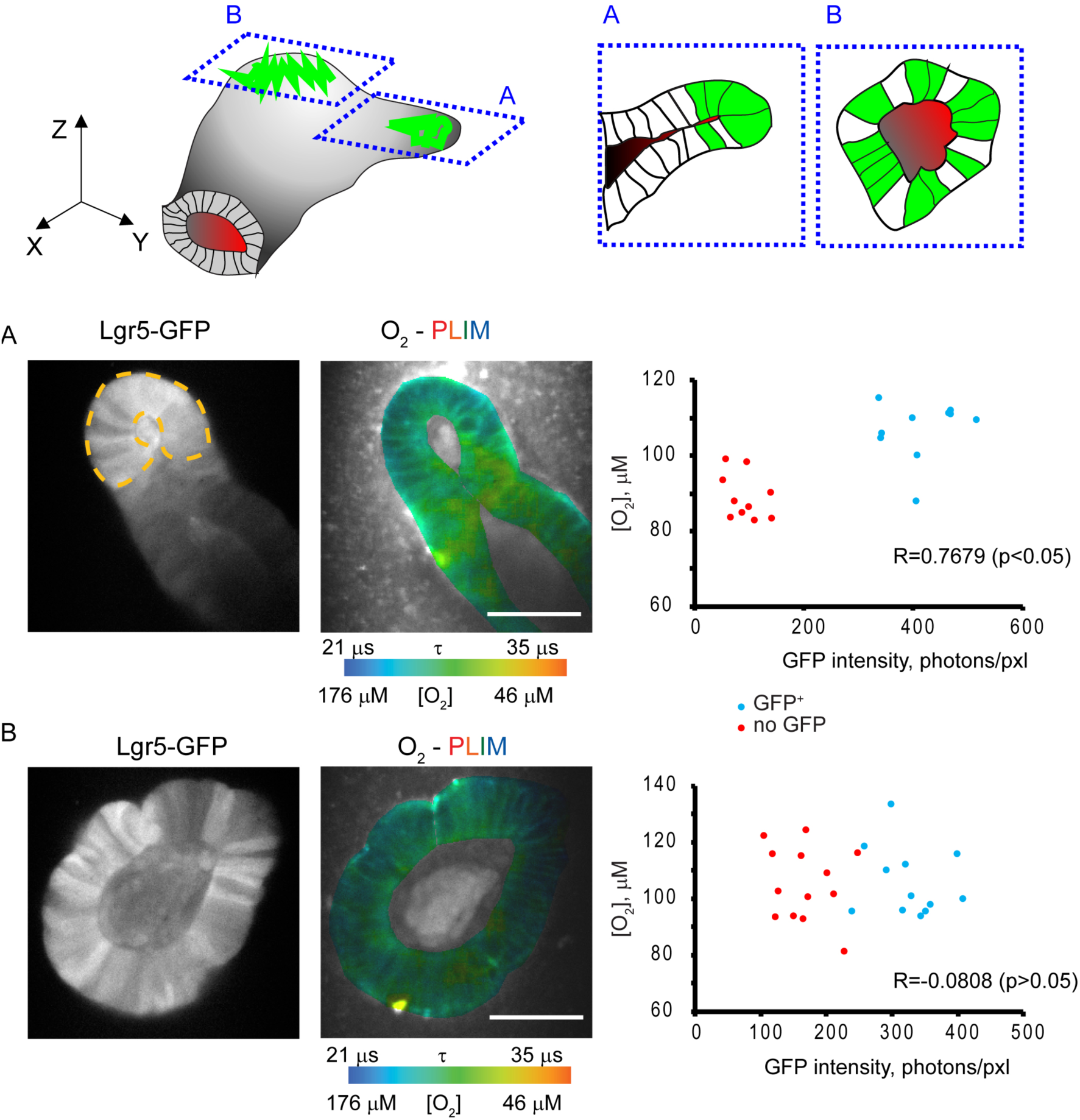
Effect of morphology and selection of Lgr5-GFP crypt regions on measured O_2_ in XY cross-sections. Top: schematic representation of intestinal organoid structure with two differently oriented regions with high expression of Lgr5-GFP, rougly corresponding to A and B images below. A, B: Lgr5-GFP intensity (grayscale) and O_2_-PLIM images with the Pearson correlation analysis for multiple ROIs within GFP^+^ (blue) and no GFP (red dots) individual cells. An ‘ideal’ cross section in A displays clear difference in oxygenation (R is close to 1), while simply different ‘mosaic’ cross-section B masks these differences (R<0). Scale bar is 50 μm.

Another factor that can confound the analysis of intestinal organoid oxygenation is non-ideal behavior of Lgr5-GFP as a label: GFP signals do not disappear immediately after cells cease proliferation and begin differentiation. Thus, the functionally heterogeneous population of GFP^+^cells represents true stem cells and the actively proliferating daughter cells being at different stages of cell cycle and differentiation stage. We also noticed that some organoids could spontaneously lose GFP labeling under ENRVC growth conditions (data not shown).

The heterogeneity of daughter cells cell-cycle within the stem cell niche is another factor potentially affecting measurements of O_2_: it is known that mitochondrial membrane polarization changes during the cell cycle and reaches a maximum during early S phase ^49^ suggesting that stem cell niche-residing cells can have enhanced utilization of OxPhos cyclically. Such mitochondrial heterogeneity within GFP^+^cells in intestinal organoids was recently reported^18^.

Understanding the functional heterogeneity of differentiated epithelial cells within organoids is also important for an accurate interpretation of results. Due to limited availability of live imaging cell-specific tracers, we did not label subpopulations of mature enterocytes, goblet, enteroendocrine, Paneth and other cell types. Thus, in presented form, the O_2_-PLIM method is expected to provide strong heterogeneity in ‘no GFP’ regions. A potential solution to address this is combining O_2_-PLIM, with NAD(P)H-FLIM, labeling of membrane polarization and proliferating cells in a 3D multi-parameter readout.

The assumption that the main consumer of oxygen within tissue is the activity of electron transport chain cannot be always accepted. Within the intestine, other ‘respiration’ processes can cause apparent oxygenation heterogeneity, for example the enterocyte-specific expression of cytochromes P450 (CYP450) ^50^. On the other hand, the heterogeneity of reactive oxygen species production and inactivation between the different cell types in the intestine is rarely considered ^5, 51^.

The application of microbiota into the lumen was not investigated in this study but seemingly intestinal epithelium by itself can sustain strong physiological (de)oxygenation, in agreement with recent *in vivo* data ^52^.

### Dynamics of organoid metabolism revealed by FLIM and O_2_-PLIM

PLIM enables direct measurement of O_2_ – a marker of the activity of electron transport chain and by other processes (non-mitochondrial respiration). NAD(P)H-FLIM detects alteration in NAD(P)H levels, which can significantly vary within the intestinal epithelium. Although this method does not give direct information on electron transport chain activity, data on NAD(P)H upon different metabolic treatments are highly useful for analysis of combined pools of NADH and NADPH. Indeed, employing both of these methods enables deeper understanding of organoid metabolism. The summary of the methodologies is present in Figure 6.

**Figure 6.**
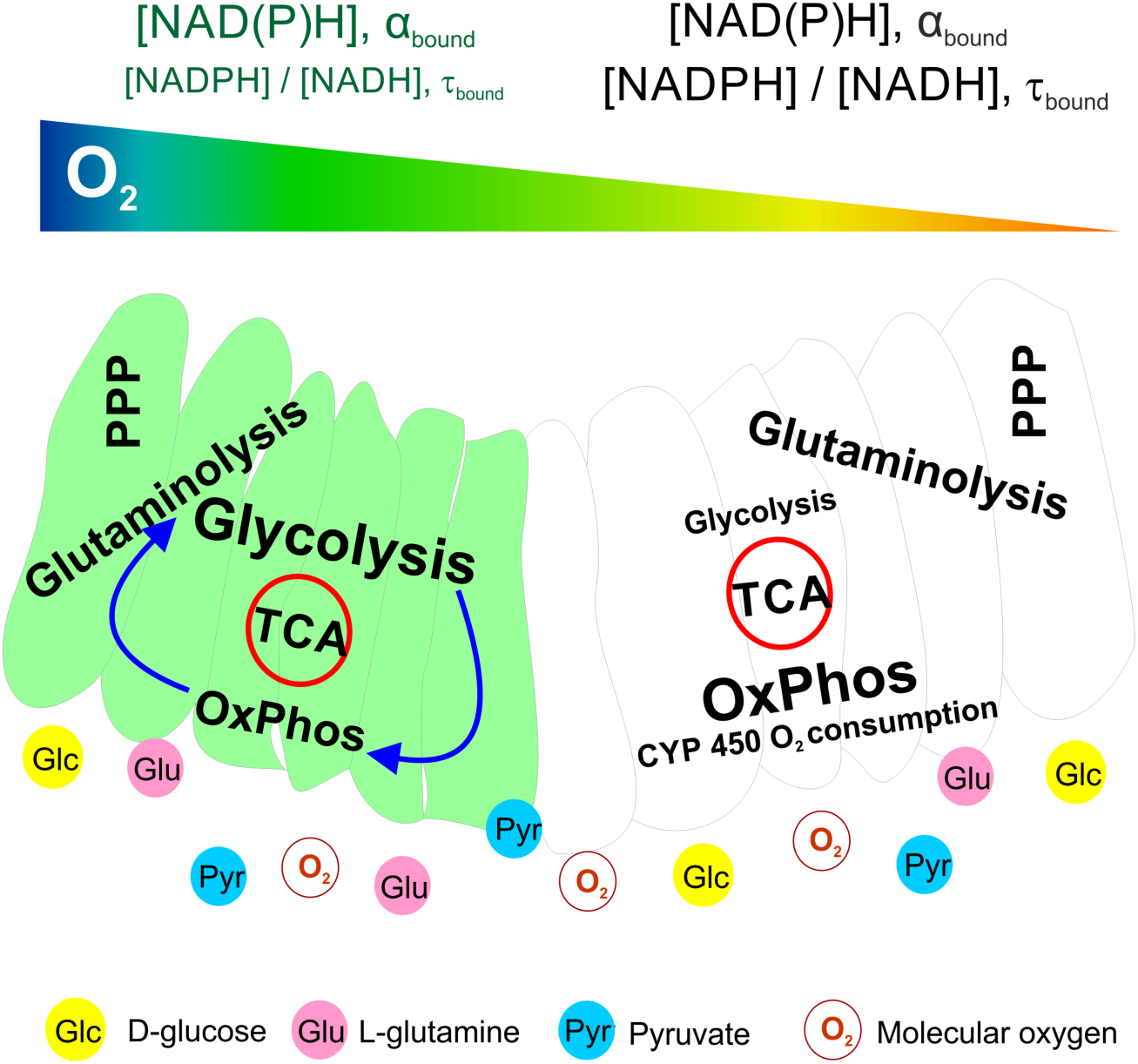
Schematic representation of the intestinal organoid cell metabolism in Lgr5-GFP^+^ cells (green) and differentiated cells. TCA – tricarboxylic acid cycle; PPP – pentose phosphate pathway; CYP450 – intestinal cytochromes P450.

Thus, by using combination of these methods, we found that O_2_ heterogeneity in the intestinal organoids has functional meaning, caused by the differences in steady-state O_2_ between the stem cell niche (Lgr5-GFP^+^ crypt regions) and mature cells (no GFP-expressing villi cells). Strikingly, oxidative metabolism levels in organoids was active, in that we saw relatively high resting deoxygenation corresponding to 7-9% O_2_. We demonstrated that on average GFP^+^regions had higher oxygenation levels (reflecting their higher reliance on glycolysis) than more actively respiring no GFP regions. However due to current limitations of available instrumentation, the O_2_-PLIM method was not sensitive enough to detect changes in oxygenation under varying media conditions (see ‘Methodological limitations’ above). Regardless the variability of oxygenation, O_2_-PLIM displayed trending increase of O_2_ difference between GFP^+^ and no GFP regions in low glucose conditions. This phenomenon can be explained by an increasing mitochondrial respiration in both organoid regions (GFP^+^and no GFP) and additional respiration processes such as activity of cytochrome P450 in the enterocytes. The decrease of this oxygenation difference in no pyruvate media (Fig. 2) can be explained by a diminishing electron transport chain activity in mature no GFP areas, highly dependent on pyruvate-fueled production of NADH, in agreement with our NAD(P)H-FLIM results (Fig. 4). We found that GFP^+^regions demonstrated ‘flexible’ balance of energy production pathways to support production of NADH by Krebs cycle on appropriate levels and to keep the functional electron transport chain and OxPhos (Fig. 6). Potentially, future comparison between LG and no PYR groups can help detecting significant changes in their oxygenation differences.

NAD(P)H-FLIM of GFP^+^and no GFP regions confirmed their principally different response to decrease of glucose and pyruvate. Our results indicate that for no GFP mature regions it is of higher priority to sustain [NADPH]/[NADH] ratio even if it would decrease the total NAD(P)H pool, while stem GFP^+^cells require a constant pool of protein-bound NAD(P)H. This can be explained by different requirements in NADPH within the proliferating stem cell niche and differentiated cells (consisting of mostly enterocytes and rare population of enteroendocrine and goblet cells) due to excess glucose present in the organoid microenvironment, an inefficient but quicker metabolic production of ATP might be optimal for high energetic demands ^53^. Since the fraction of bound NAD(P)H pool in ‘no GFP’ areas was sensitive to pyruvate decrease, the key metabolite for acetyl-CoA synthesis and fuel of Krebs cycle, the Krebs cycle and OxPhos (main sources of ATP, NADH and intermediate metabolites) must be very active in differentiated cells (Fig. 6). In addition, it has been shown that differentiation of intestinal stem cell results in an increase of the mitochondrial pyruvate carrier gene expression; this renders differentiated cells more dependent on the presence of endogenous and exogenous pyruvate pools, while stem cells are rather unaffected ^6^. The presence of high glucose during the pyruvate depletion experiment did not compensate the effect on the NAD(P)H bound fraction, suggesting that glycolysis has generally low rate in no GFP cells. This explains the observed decrease of differences in oxygenation between GFP^+^and no GFP regions in no-PYR media (glycolysis is not efficient to fuel OxPhos) in comparison to PYR (HG) media (Fig. 2H). While glutaminolysis can be used to sustain Krebs cycle ^54-56^, the observed ability to keep high [NADPH]/[NADH] ratio in ‘no GFP’ cells could be due to the high activity of pentose phosphate pathway, mitochondrial nicotinamide nucleotide transhydrogenase and malic enzymes as main producers of NADPH in cells ^28, 57, 58^. Indeed the glucose-dependent NADPH production is important for activity of enterocyte glutathione reductase, the principal supplier of GSSH in the intestine ^59^. Activities of intestinal enterocyte-specific enzymes such as cytochrome P450 oxidase, reductase and NADPH oxidase 4 (NOX4) also demand for high levels of NADPH ^31, 60, 61^.

GFP^+^stem cell niche regions displayed [NADPH]/[NADH] ratio sensitive to presence of glucose and pyruvate, but their protein-bound NAD(P)H fraction tended to remain constant (Fig. 4C, D). This confirms their high glycolytic metabolism, which can be easily switched to OxPhos when necessary. We believe that such adaptability facilitates stem cells to sustain levels of NAD(P)H and intermediate metabolites to maintain ATP production at levels supporting cell proliferation. The observed increase of bound [NADPH]/[NADH] ratio in low glucose media can be functionally important as NADPH is directly involved in protection from reactive oxygen species produced due to electron leaks within the electron transport chain.

Our findings are also in agreement with observations of Stringari and co-workers ^32^ even though this study dealt with *ex vivo* intestinal tissue exposed to conditions favoring cell starvation, potential hypoxia in crypt regions and did not take into account the NADPH.

In conclusion, this present and previous studies emphasize the importance of viewing cell fate regulation in organoids and regenerating tissues from the point of growth media composition and nutrient availability. Emerging live imaging microscopy of cell metabolism is of high relevance to this emerging field of interest. The applied O_2_-PLIM and NAD(P)H-FLIM methods enable visualization of metabolic discrepancies in stem cell niche and differentiated cells corresponding to crypt and villi compartments of the small intestinal epithelium. Future application of these methods together in multi-parametric imaging with appropriate cell type markers will allow the study of organoid stem cell metabolism with single cell and subcellular resolution.

## Methods

### Materials

N2 Supplement, 100x concentrate (17502-048), B27 media Supplement, 50x concentrate (17504-044) and GlutaMax Supplement 100x concentrate (35050-038) were acquired from Invitrogen (GE Healthcare, Ireland). Recombinant murine EGF (epidermal growth factor) (315-09), recombinant human R-spondin-1 (120-38) and recombinant murine Noggin (250-38) were from Peprotech (NJ, USA). Matrigel^®^ with reduced growth factors (356231) was from Corning (VWR, Ireland). Penicillin-streptomycin solution (P0781), *N*-Acetyl-l-cysteine (NAC, A9165), Dulbecco’s modified Eagle’s medium nutrient mixture F-12 [HAM] (D6421), Dulbecco’s modified Eagle’s medium (DMEM), phenol red-, glucose-, pyruvate- and glutamine-free (D5030), D(+)-Glucose (G8270), 1 M HEPES solution pH 7.2 (H0887), 100 mM sodium pyruvate (S8636), 200 mM L-Glutamine (G7513), CHIR99021 (SML1046) and sodium valproate (P4543) and all the other reagents were from Sigma-Aldrich (Dublin, Ireland). Phosphorescent O_2_-sensitive Pt-Glc probe was synthesized as described previously ^17^. Seahorse XF96 plates were from Agilent^®^ technologies (Little Island, Cork, Ireland).

### Lgr5-GFP organoid culture and staining with O_2_ probe

Lgr5-GFP (*Lgr5-EGFP-ires-CreERT2*) mouse intestinal organoid culture^35^ were kindly provided by Prof. H. Clevers Lab (Hubrecht Institute, The Netherlands). Organoids were handled as described previously^18^ and typically were grown in ENRVC media, consisting of Dulbecco modified Eagle’s medium nutrient mixture F-12 [HAM] supplemented with 1% penicillin-streptomycin, 1% GlutaMax, 1% N2 supplement, 2% B-27 supplement, 1.25 mM NAC, 50 ng/ml EGF, 1 μg/ml R-spondin-1, 100 ng/ml Noggin, 1 mM sodium valproate and 3 μM CHIR99021. Intact organoids were seeded onto 20 μl of Matrigel in 35 mm tissue culture treated dishes (Sarstedt, Dublin, Ireland) in ENRVC medium, 1-2 days prior to imaging. Staining with O_2_-probe was achieved by incubating organoids with 2 μM Pt-Glc diluted either in a Phenol red-free DMEM supplemented with 10 mM HEPES pH 7.2, 2 mM glutamine, 10 mM D-glucose, 1 mM sodium pyruvate (HG), the same medium with 0.5 mM D-glucose (LG) or with no pyruvate (no-PYR) for 1 h, followed by washing in respective media. Imaging was performed during 30 min after staining. Two experimental replicates for each media combinations (e.g. HG vs. LG, HG vs. no-PYR) were performed on separate cultures originated from different frozen organoid aliquots.

### Analysis of oxygen consumption rates of organoids on a microplate

This was performed accordingly to published protocol ^19^, with minor modifications. 9-10 organoids in 3 μl of Matrigel^®^ per well were seeded onto XF96 plate (Seahorse) and subsequently grown in ENRVC or EN (lacking R-spondin 1, sodium valproate and CHIR99021) media for 2 days prior to the analysis. The number of organoids per well was calculated manually and used for normalization.

### O_2_-PLIM and data analysis

An upright Zeiss Axio Examiner Z1 microscope equipped with water dipping 63x/1.0 W-Plan Apochromat dipping objective, DCS-120 confocal scanner (Becker & Hickl GmbH), heated incubator and stage (t=37 °C), motorized Z-axis control and 405 nm and 408 nm BDL-SMNI pulsed diode lasers (Becker & Hickl GmbH) was used for O_2_-PLIM. Emission was collected using 635-675 nm (405 nm excitation, O_2_ probe) and 512-536 nm (488 nm excitation, Lgr5-GFP) bandpass filters, essentially as described previously using 256×256 XY scanning resolution ^16, 17^. Each imaged optical section corresponding to individual GFP^+^region was considered as an experimental unit in the statistical analysis. Combined analysis (segmentation, fitting of phosphorescence decays) of Lgr5-GFP fluorescence and O_2_-PLIM images was achieved using SPCImage software (Becker & Hickl GmbH, Germany). Mono-exponential fit (with χ^2^ <1.5) was used on a pixel-by-pixel base to calculate phosphorescence lifetimes and produce color-coded oxygen distribution images. Lgr5-GFP intensity (photons), Pt-Glc intensity (photons) and phosphorescence lifetime (color-coded values) data were exported and processed in Microsoft Excel, as described previously^43^. For analysis, we used ROIs with size of 5×5 Excel table (∼4×4 μm size). Since the GFP intensity was highly variable, we limited our analysis to the regions, where GFP intensity was comparable or higher than the luminal autofluorescence. ‘No GFP’ ROIs were considered when their intensity was at least 3 times lower than for neighbor GFP^+^cells.

Selected 9-12 ROIs with calculated O_2_ for each experimental group (GFP^+^, no GFP) were analyzed with the Kolmogorov-Smirnov (K-S) test for normality and used for calculation of average values and differences in oxygenation between (GFP^+^and no GFP). A Pearson correlation analysis (between GFP intensity and oxygenation) and statistical analysis of oxygenation with independent *t*-test (at p = 0.1 or p = 0.05) was applied to each optical section. The comparison between GFP^+^and no GFP oxygenation in separate groups was also done with paired *t*-test in Origin 6.0 software.

### Two-photon excited NAD(P)H-FLIM

This was performed using a custom upright (Olympus BX61WI) laser multiphoton microscopy system equipped with Titanium:sapphire laser (Chameleon Ultra, Coherent^®^, USA) and water-immersion 25x objective (Olympus, 1.05NA) and temperature controlled stage. Two-photon excitation of NAD(P)H and GFP fluorescence was performed at excitation wavelengths of 760 nm and 920 nm, respectively. A 455/90 nm bandpass filter was used to isolate the NAD(P)H fluorescence emission and a 502/47 nm bandpass filter for GFP fluorescence emission. 512 × 512 pixel images were acquired with a pixel dwell time of 3.81 μs and 30-second collection time. A PicoHarp 300 TCSPC system operating in the time-tagged mode coupled with a photomultiplier detector assembly (PMA) hybrid detector (PicoQuanT GmbH, Germany) was used for fluorescence decay measurements yielding 256 time bins per pixel. For each intestinal crypt, a GFP FLIM image was acquired followed by the acquisition of a NAD(P)H FLIM image. After GFP FLIM imaging, the fluorescence decay was measured allowing identifying GFP^+^and no GFP areas confirmed by the measurement of GFP fluorescence lifetime. The same regions were overlapped in the NAD(P)H FLIM images to perform a fluorescence decay analysis.

Intestinal organoids were pre-incubated (2-3 h) in respective HG, LG or no-PYR media and least 11 crypts from each treatment group were analyzed.

Fluorescence lifetime images with their associated decay curves for NAD(P)H and GFP were obtained. The overall decay curve was generated by the contribution of all pixels and was fitted with a mono- or a double exponential decay curve, for GFP and NAD(P)H, respectively (Eq. 1)

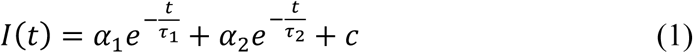

*I(t)* corresponds to the fluorescence intensity measured at time *t* after laser excitation; *α1* and *α*_*2*_ represents the fraction of the overall signal proportion of a short and long component lifetime components, respectively. For a mono-exponential decay, *α*_*2*_ is zero, making *a*_*1*_ a fraction corresponding to 100% of the overall fluorescence signal. *τ*_*1*_ and *τ*_*2*_ are the short and long lifetime components, respectively; *C* corresponds to background light. χ^2^ represents the goodness of multi-exponential fit to the raw fluorescence decay data. In this study all the values with χ^2^ < 1.3 were considered as ‘good’ fits.

For NAD(P)H, a two-component fitting was also used to differentiate between the free (*τ*_*1*_) and protein-bound (*τ*_*2*_) NAD(P)H: the average lifetime (*τ*_ave_) of NAD(P)H for each pixel was calculated by a weighted average of both free and bound lifetime contributions (Eq. 2).

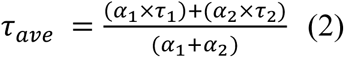

## Statistical Analysis

The data produced by different methodologies were assessed and evaluated for statistical significance as described in respective sections. Briefly, results were tested for normality of distribution, using K-S test, with subsequent analysis by independent or paired (where appropriate) *t*-test. NAD(P)H-FLIM and O_2_-PLIM data were analyzed separately, by different individuals. Details of used tests are provided in Supplementary tables 1 and 2 and in respective figure legends.

## Acknowledgments

We thank Dr. Jens Puschoff and Prof. Hans Clevers (Hubrecht Institute) for support with Lgr5-GFP organoids and Dr. Ryan McGarrigle (Agilent Technologies) for help with XF96 microplate analysis.

This work was supported by the Agilent University Research Program (ACT-UR, No. 4225), Science Foundation Ireland (SFI) grants 13/SIRG/2144 and 18/IF/6238 and the Russian Science Foundation (RSF, 18-15-00407). NN is supported by a Trinity College Dublin, Provost’s Project PhD Award and the TCD FLIM Core unit directed by MM is supported by a SFI Infrastructure Programme: Category D Opportunistic Funds Call.

## Supplementary Tables

**Table S1.** Summary of produced O_2_ imaging data between different experiments. Significantly different (p<0.1) data are indicated with shading. HG-high glucose medium (equals to PYR, 10 mM glucose), LG – low glucose medium (0.5 mM glucose); NP – no pyruvate medium (10 mM glucose).

| Replicate | Treatment | Average [O<sub>2</sub>] ± stdev (N), μM |  |  |  | Average gradient d[O<sub>2</sub>] ± stdev (N), μM |  |
| --- | --- | --- | --- | --- | --- | --- | --- |
|  |  | whole organoids |  | GFP<sup>+</sup> areas | GFP<sup>-</sup> areas |  |  |
| 1-1 | HG (10 mM d-glucose, 1 mM pyruvate Na) | 71±19 (14) | Independent t-Test<br>p = 0.972 | 66±21 (8) | 63±20 (8) | 2.5±4.5 (8) | Independent t-Test<br>p = 0.12 |
|  |  |  |  | Paired t-Test<br>p = 0.164 |  |  |  |
|  | LG (0.5 mM d-glucose, 1 mM pyruvate Na) | 71±13 (13) |  | 73±15 (11) | 67±13 (11) | 5.8±5.9 (11) |  |
|  |  |  |  | Paired t-Test<br>p = 0.009 |  |  |  |
| 1-2 | HG | 89±21 (15) | Independent t-Test<br>p = 0.685 | 91±21 (15) | 86±22 (15) | 4.5±8.9 (15) | Independent t-Test<br>p = 0.463 |
|  |  |  |  | Paired t-Test<br>p = 0.068 |  |  |  |
|  | LG | 92±18 (15) |  | 95±18 (15) | 88±18 (15) | 6.5±4.7 (15) |  |
|  |  |  |  | Paired t-Test<br>p =0.0001 |  |  |  |
| 1-1 and 1-2 | HG | 80±22 (29) | Independent t-Test<br>p = 0.743 | 82±24 (23) | 78±24 (23) | 3.8±7.6 (23) | Independent t-Test<br>p = 0.204 |
|  |  |  |  | Paired t-Test<br>p = 0.025 |  |  |  |
|  | LG | 82±19 (28) |  | 85±20 (26) | 79±19 (26) | 6.2±5.1 (26) |  |
|  |  |  |  | Paired t-Test<br>p = 0.000002 |  |  |  |
| 2-1 | NP (10 mM d-glucose, no sodium pyruvate) | 70±26 (9) | Independent t-Test<br>p = 0.545 | 85±27 (5) | 81±25 (5) | 4±4.6 (5) | Independent t-Test<br>p = 0.483 |
|  |  |  |  | Paired t-Test<br>p = 0.122 |  |  |  |
|  | PYR (equivalent to HG medium) | 75±12 (11) |  | 79±12 (11) | 72±13 (11) | 6.6±7.3 (11) |  |
|  |  |  |  | Paired t-Test<br>p = 0.013 |  |  |  |
| 2-2 | NP | 80±15 (12) | Independent t-Test<br>p = 0.492 | 82±15 (11) | 77±15 (11) | 5.7±4.9 (11) | Independent t-Test<br>p = 0.442 |
|  |  |  |  | Paired t-Test<br>p = 0.003 |  |  |  |
|  | PYR | 76±13<br>(12) |  | 80±14<br>(11) | 72±11<br>(11) | 8±8.8<br>(11) |  |
|  |  |  |  | Paired t-Test<br>p = 0.012 |  |  |  |
| 2-1 and<br>2-2 | NP | 76±20<br>(21) | Independent<br>t-test<br>p = 0.999 | 83±19<br>(16) | 78±18<br>(16) | 5.1±4.7<br>(16) | Independent<br>t-test<br>p = 0.336 |
|  |  |  |  | Paired t-Test<br>p = 0.0006 |  |  |  |
|  | PYR | 76±12<br>(23) |  | 79±13<br>(22) | 72±12<br>(22) | 7.3±7.9<br>(22) |  |
|  |  |  |  | Paired t-Test<br>p = 0.0003 |  |  |  |

**Table S2.** Summary of produced NAD(P)H-FLIM imaging data between different experiments. Significantly different (p<0.1) data are indicated with shading. HG: high glucose medium (equals to PYR, 10 mM glucose), LG: low glucose medium (0.5 mM glucose); NP:no pyruvate medium (10 mM glucose).

| Replicate | Treatment | Average lifetime NAD(P)H<br>± stdev (N), ns |  | Protein-bound NAD(P)H<br>lifetime ± stdev (N), ns |  | Fraction of bound NAD(P)H<br>± stdev (N) |  |
| --- | --- | --- | --- | --- | --- | --- | --- |
|  |  | GFP<sup>+</sup> areas | GFP<sup>-</sup> areas | GFP<sup>+</sup> areas | GFP<sup>-</sup> areas | GFP<sup>+</sup> areas | GFP<sup>-</sup> areas |
| 1-1 | HG (10 mM<br>d-glucose, 1<br>mM pyruvate<br>Na) | 1.73±0.08<br>(14) | 1.82±0.12<br>(14) | 3.28±0.11<br>(14) | 3.42±0.16<br>(14) | 0.42±0.03<br>(14) | 0.42±0.04<br>(14) |
|  |  | Paired t-Test<br>p = 0.0007 |  | Paired t-Test<br>p = 0.0130 |  | Paired t-Test<br>p = 0.2817 |  |
| 1-2 | LG (0.5 mM<br>d-glucose, 1<br>mM pyruvate<br>Na) | 1.82±0.16<br>(13) | 1.81±0.13<br>(13) | 3.39±0.21<br>(13) | 3.45±0.16<br>(13) | 0.43±0.04<br>(13) | 0.41±0.03<br>(13) |
|  |  | Paired t-Test<br>p = 0.8144 |  | Paired t-Test<br>p = 0.4347 |  | Paired t-Test<br>p = 0.1693 |  |
| 1-3 | NP (10 mM<br>d-glucose, no<br>sodium<br>pyruvate) | 1.69±0.09<br>(11) | 1.73±0.10<br>(11) | 3.18±0.09<br>(11) | 3.44±0.14<br>(11) | 0.41±0.03<br>(11) | 0.38±0.03<br>(11) |
|  |  | Paired t-Test<br>p = 0.0477 |  | Paired t-Test<br>p < 0.0001 |  | Paired t-Test<br>p < 0.0001 |  |

## Supplementary figures

**Figure S1.**
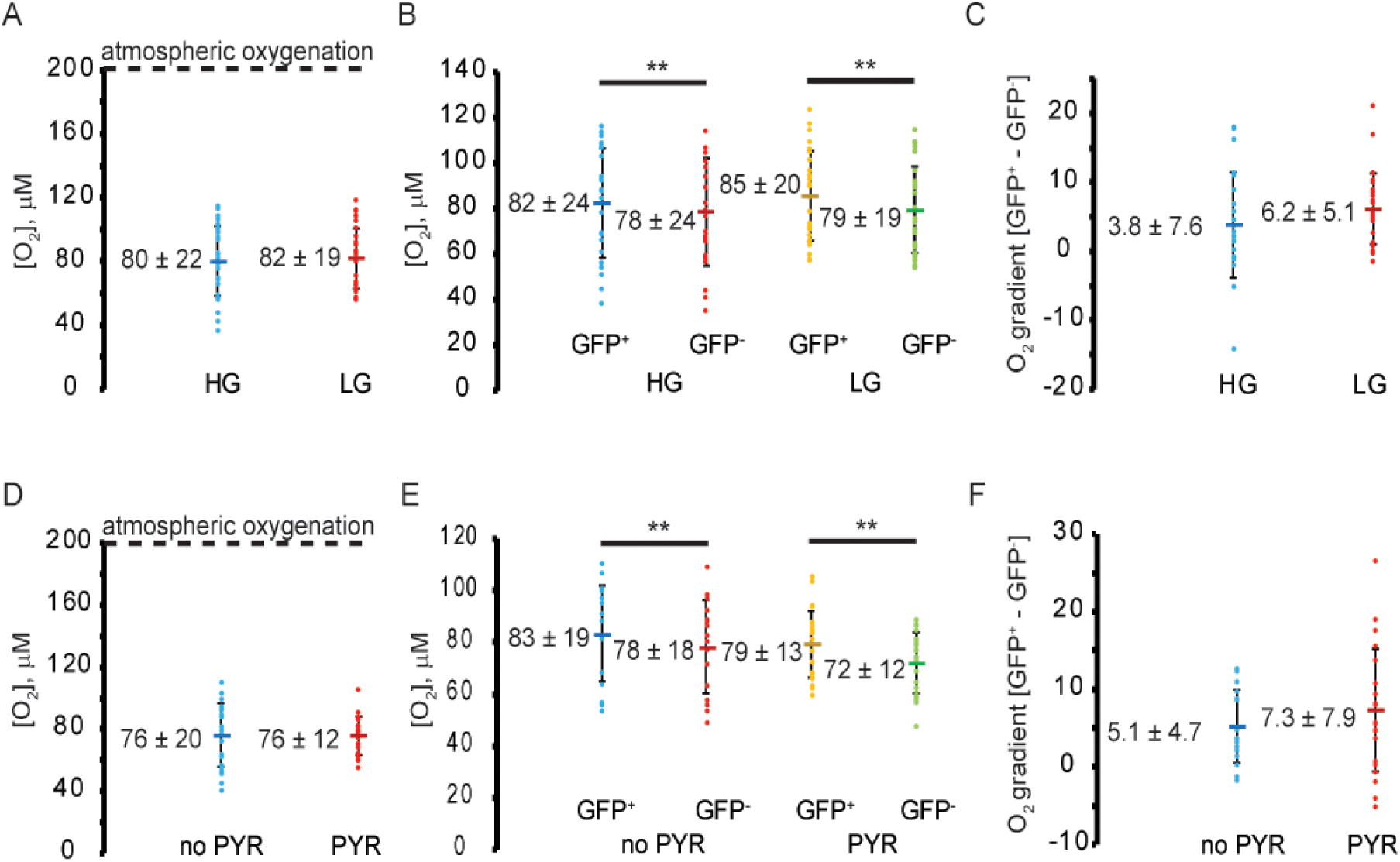
O_2_-PLIM analysis of Lgr5-GFP organoids from combined experimental repeats. Each point indicates average values calculated from the individual optical sections. Line marker indicates mean values of each group. Asterisks indicate significant differences ** p<0.05. A, D: Effect of medium composition on average oxygenation. No significant differences were found with independent *t*-test (at p=0.1). B, E: Comparison of average oxygenation between GFP^+^ and no GFP areas for different media. Paired *t*-test revealed significant differences in their average oxygenation for each media composition. No significant differences were detected between average oxygenation of GFP^+^ and no GFP areas using independent *t*-test (at p=0.1). C, F: Effect of media composition on average oxygenation difference between GFP^+^ and no GFP areas No significant differences were detected with independent *t-*test (p=0.1).

**Figure S2.**
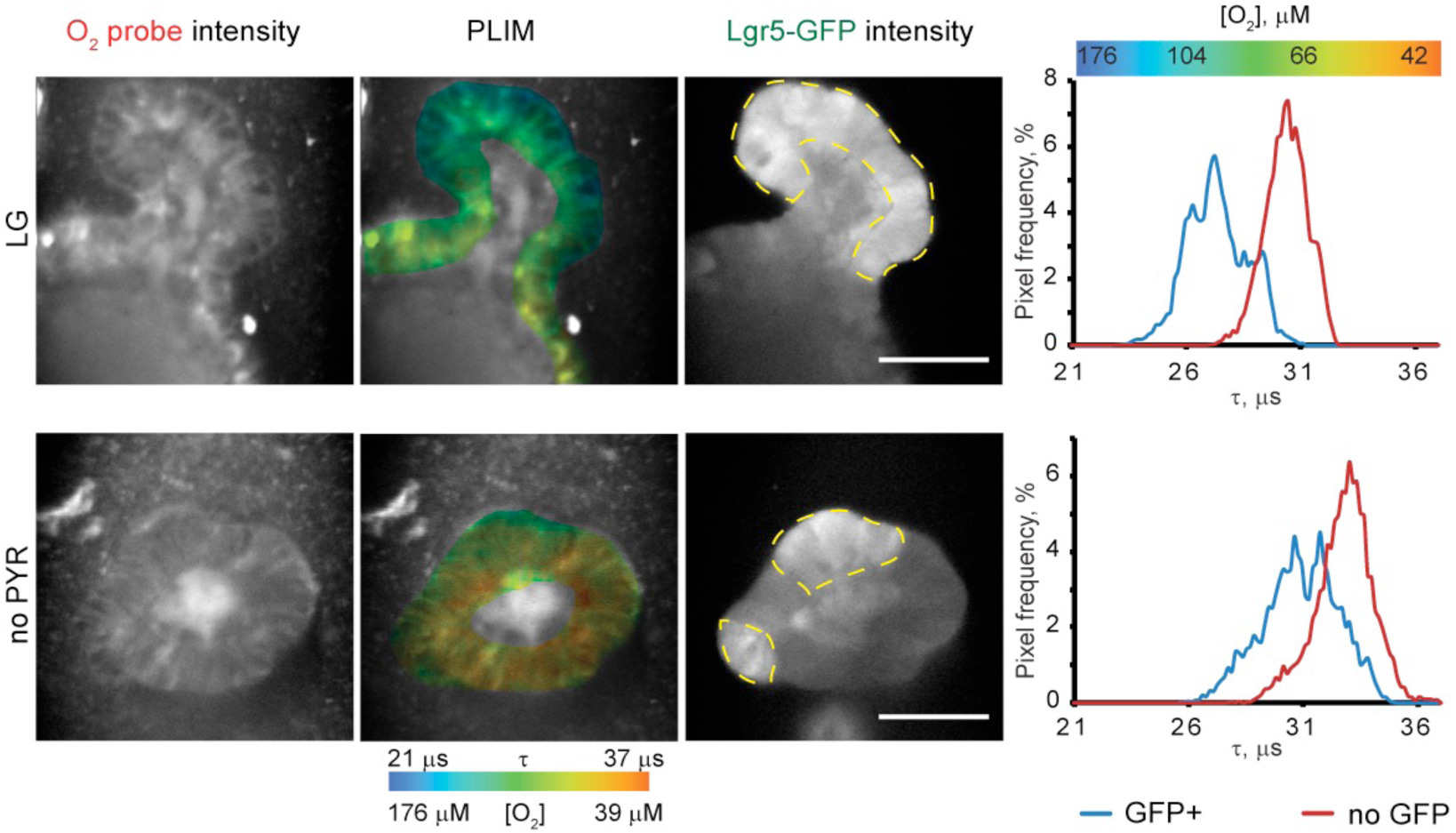
Examples of optical sections with statistically significant oxygenation differences between GFP^+^ and no GFP regions for LG and no-PYR media. Intensity, combined PLIM and GFP fluorescence images and fluorescence lifetime distribution histograms (right) are shown. Significant difference was observed with independent *t*-test of GFP^+^ and no GFP regions from individual images of intestinal organoids (at p<0.01). Oxygenation mean values ± standard deviation of GFP^+^ / no GFP areas were: 96±11 μM / 75 ±11 μM (N=9) for LG and 68±5 μM / 56±5 μM (N=9) for no PYR-treated organoids. Scale bar is 50 μm.

**Figure S3.**
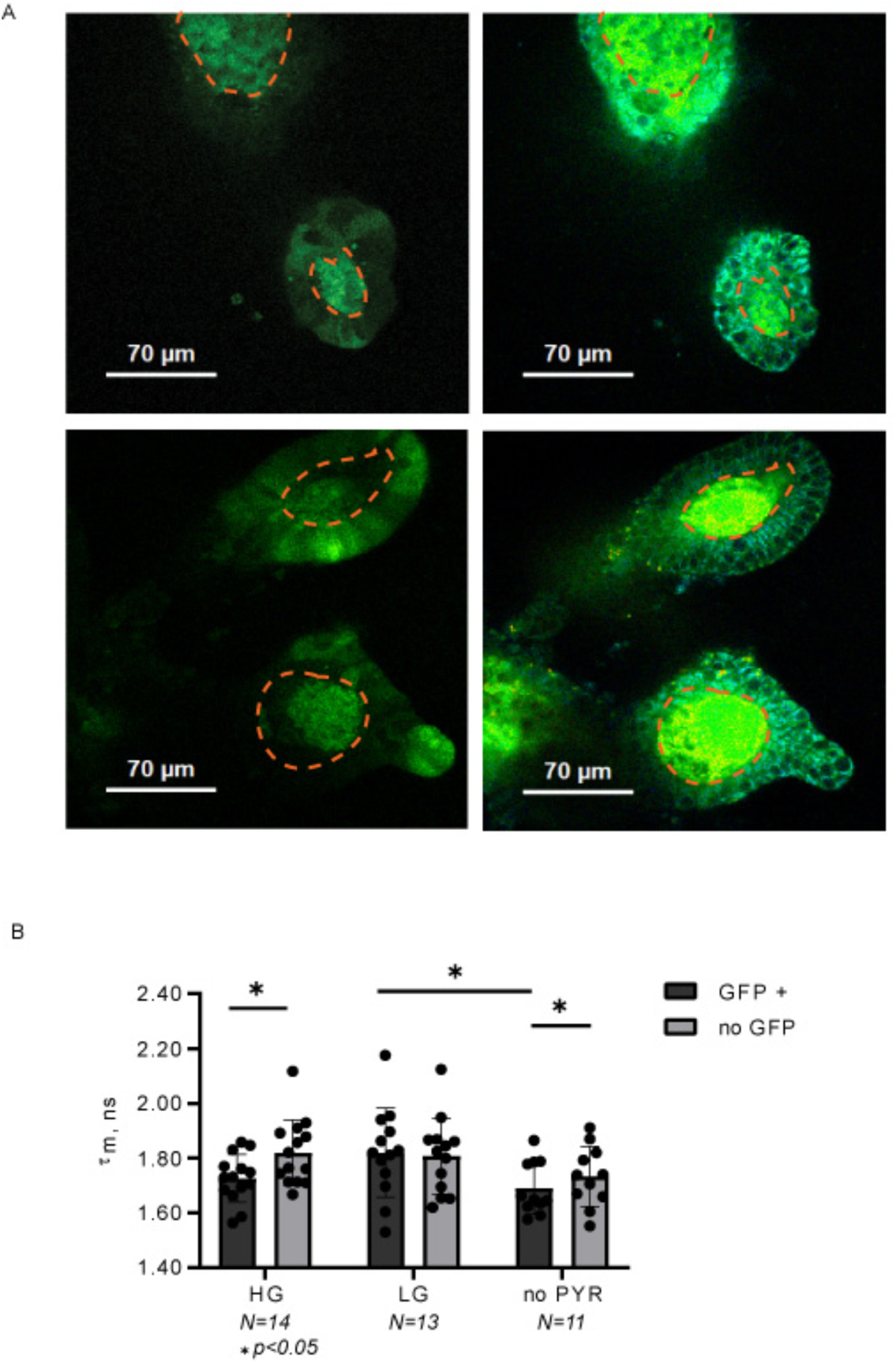
A: examples of images of Lgr5-GFP fluorescence intensity and two-photon NAD(P)H-FLIM of intestinal organoids. Lumen is indicated with a dashed line. B: summary of τmean of NAD(P)H autofluorescence measured for HG, LG and no-PYR media. Significant differences (paired *t*-test, p<0.05) are indicated with asterisks.

## References

1. Kretzschmar, K. & Clevers, H. Organoids: modeling development and the stem cell niche in a dish. Developmental cell 38, 590–600 (2016).

2. Rath, E., Moschetta, A. & Haller, D. Mitochondrial function—gatekeeper of intestinal epithelial cell homeostasis. Nature Reviews Gastroenterology & Hepatology 15, 497 (2018).

3. Shyh-Chang, N., Daley, G.Q. & Cantley, L.C. Stem cell metabolism in tissue development and aging. Development 140, 2535–2547 (2013).

4. Oh, J., Lee, Y.D. & Wagers, A.J. Stem cell aging: mechanisms, regulators and therapeutic opportunities. Nature medicine 20, 870 (2014).

5. Rodríguez-Colman, M.J. et al. Interplay between metabolic identities in the intestinal crypt supports stem cell function. Nature 543, 424 (2017).

6. Schell, J.C. et al. Control of intestinal stem cell function and proliferation by mitochondrial pyruvate metabolism. Nature Cell Biology 19, 1027 (2017).

7. Kumar, N. et al. A YY1-dependent increase in aerobic metabolism is indispensable for intestinal organogenesis. Development 143, 3711–3722 (2016).

8. Srivillibhuthur, M. et al. TFAM is required for maturation of the fetal and adult intestinal epithelium. Developmental biology 439, 92–101 (2018).

9. Zwick, R.K., Ohlstein, B. & Klein, O.D. Intestinal renewal across the animal kingdom: comparing stem cell activity in mouse and Drosophila. American Journal of Physiology-Gastrointestinal and Liver Physiology 316, G313–G322 (2018).

10. Clevers, H. Modeling Development and Disease with Organoids. Cell 165, 1586–1597 (2016).

11. Alonso, S. & Yilmaz, Ö.H. Nutritional regulation of intestinal stem cells. Annual review of nutrition 38, 273–301 (2018).

12. Lancaster, M.A. & Knoblich, J.A. Organogenesis in a dish: modeling development and disease using organoid technologies. Science 345, 1247125 (2014).

13. Leushacke, M. & Barker, N. Ex vivo culture of the intestinal epithelium: strategies and applications. Gut 63, 1345–1354 (2014).

14. Lyons, J., Herring, C.A., Banerjee, A., Simmons, A.J. & Lau, K.S. Multiscale analysis of the murine intestine for modeling human diseases. Integrative Biology 7, 740–757 (2015).

15. Rios, A.C. & Clevers, H. Imaging organoids: a bright future ahead. Nature Methods 15, 24 (2018).

16. Okkelman, I.A., Foley, T., Papkovsky, D.B. & Dmitriev, R.I. Live cell imaging of mouse intestinal organoids reveals heterogeneity in their oxygenation. Biomaterials 146, 86–96 (2017).

17. Okkelman, I.A., Foley, T., Papkovsky, D.B. & Dmitriev, R.I. Multi-Parametric Imaging of Hypoxia and Cell Cycle in Intestinal Organoid Culture, in Multi-Parametric Live Cell Microscopy of 3D Tissue Models, Vol. 1035. (ed. R. Dmitriev) 85-103 (Advances in Experimental Medicine and Biology, Springer, Cham; 2017).

18. Okkelman, I.A., Papkovsky, D.B. & Dmitriev, R.I. Estimation of the mitochondrial membrane potential using fluorescence lifetime imaging microscopy. Cytometry Part A: The Journal of the International Society for Analytical Cytology, in press (2019).

19. Bas, T. & Augenlicht, L.H. Real time analysis of metabolic profile in ex vivo mouse intestinal crypt organoid cultures. JoVE (Journal of Visualized Experiments), e52026 (2014).

20. Fan, Y.-Y. et al. A bioassay to measure energy metabolism in mouse colonic crypts, organoids, and sorted stem cells. American Journal of Physiology-Gastrointestinal and Liver Physiology 309, G1–G9 (2015).

21. MacDonald, J.A. et al. A nanoscale, multi-parametric flow cytometry-based platform to study mitochondrial heterogeneity and mitochondrial DNA dynamics. Communications biology 2 (2019).

22. Dmitriev, R.I. Multi-Parametric Live Cell Microscopy of 3D Tissue Models, Vol. 1035. (Springer, 2017).

23. Berezin, M.Y. & Achilefu, S. Fluorescence lifetime measurements and biological imaging. Chemical reviews 110, 2641–2684 (2010).

24. Sarder, P., Maji, D. & Achilefu, S. Molecular probes for fluorescence lifetime imaging. Bioconjugate chemistry 26, 963–974 (2015).

25. Dmitriev, R.I., Zhdanov, A.V., Nolan, Y.M. & Papkovsky, D.B. Imaging of neurosphere oxygenation with phosphorescent probes. Biomaterials 34, 9307–9317 (2013).

26. Papkovsky, D.B. & Dmitriev, R.I. Imaging of oxygen and hypoxia in cell and tissue samples. Cellular and molecular life sciences 75, 2963–2980 (2018).

27. Ma, N., Digman, M.A., Malacrida, L. & Gratton, E. Measurements of absolute concentrations of NADH in cells using the phasor FLIM method. Biomedical optics express 7, 2441–2452 (2016).

28. Blacker, T.S. et al. Separating NADH and NADPH fluorescence in live cells and tissues using FLIM. Nature communications 5, 3936 (2014).

29. Okkelman, I.A., Dmitriev, R.I., Foley, T. & Papkovsky, D.B. Use of fluorescence lifetime imaging microscopy (FLIM) as a timer of cell cycle S phase. PloS one 11, e0167385 (2016).

30. Brand, M.D. & Nicholls, D.G. Assessing mitochondrial dysfunction in cells. Biochemical Journal 435, 297–312 (2011).

31. Lindquist, R., Bayat-Sarmadi, J., Leben, R., Niesner, R. & Hauser, A. NAD (P) H oxidase activity in the small intestine is predominantly found in enterocytes, not professional phagocytes. International journal of molecular sciences 19, 1365 (2018).

32. Stringari, C. et al. Metabolic trajectory of cellular differentiation in small intestine by Phasor Fluorescence Lifetime Microscopy of NADH. Scientific reports 2, 568 (2012).

33. Kalinina, S. et al. Correlative NAD (P) H-FLIM and oxygen sensing-PLIM for metabolic mapping. Journal of biophotonics 9, 800–811 (2016).

34. O’Donnell, N. et al. Cellulose-based scaffolds for fluorescence lifetime imaging-assisted tissue engineering. Acta biomaterialia 80, 85–96 (2018).

35. Barker, N. et al. Identification of stem cells in small intestine and colon by marker gene Lgr5. Nature 449, 1003 (2007).

36. Sato, T. et al. Single Lgr5 stem cells build crypt–villus structures in vitro without a mesenchymal niche. Nature 459, 262 (2009).

37. Dmitriev, R.I. et al. Small molecule phosphorescent probes for O 2 imaging in 3D tissue models. Biomaterials Science 2, 853–866 (2014).

38. Zhdanov, A.V., Golubeva, A.V., Okkelman, I.A., Cryan, J.F. & Papkovsky, D. Imaging of oxygen gradients in giant umbrella cells: an ex vivo PLIM study. American Journal of Physiology-Cell Physiology 309, C501–C509 (2015).

39. Zhdanov, A.V., Okkelman, I.A., Collins, F.W., Melgar, S. & Papkovsky, D.B. A novel effect of DMOG on cell metabolism: direct inhibition of mitochondrial function precedes HIF target gene expression. Biochimica et Biophysica Acta (BBA)-Bioenergetics 1847, 1254–1266 (2015).

40. Koren, K. et al. Luminescence Lifetime Imaging of Chemical Sensors – a Comparison between Time-Domain and Frequency-Domain based Camera Systems. Analytical Chemistry (2019).

41. Le Marois, A. & Suhling, K. Quantitative Live Cell FLIM Imaging in Three Dimensions, in Multi-Parametric Live Cell Microscopy of 3D Tissue Models 31–48 (Springer, 2017).

42. Sakadžić, S. et al. Two-photon high-resolution measurement of partial pressure of oxygen in cerebral vasculature and tissue. Nature methods 7, 755 (2010).

43. Okkelman, I.A., Puschhof, J., Papkovsky, D.B. & Dmitriev, R.I. Visualization of stem cell niche by the fluorescence lifetime imaging microscopy (FLIM), in Intestinal Stem Cells: methods and protocols, Vol. In press. (ed. P. Ordonez Moran) XXXX (Advances in Experimental Medicine and Biology, Springer, Cham; 2020).

44. Walsh, A.J. et al. Quantitative optical imaging of primary tumor organoid metabolism predicts drug response in breast cancer. Cancer research 74, 5184–5194 (2014).

45. Dmitriev, R.I. & Papkovsky, D.B. Intracellular probes for imaging oxygen concentration: how good are they? Methods and applications in fluorescence 3, 034001 (2015).

46. Müller, B.J. et al. Nanoparticle-Based Fluoroionophore for Analysis of Potassium Ion Dynamics in 3D Tissue Models and In Vivo. Advanced functional materials 28, 1704598 (2018).

47. Dmitriev, R.I. & Papkovsky, D.B. Quenched-phosphorescence detection of molecular oxygen: applications in life sciences, Vol. 11. (Royal Society of Chemistry, 2018).

48. Lakner, P., Monaghan, M.G., Moller, Y., Olayioye, M. & Schenke-Layland, K. Applying phasor approach analysis of multiphoton FLIM measurements to probe the metabolic activity of 3D in vitro cell culture models. Sci Rep 7, 42730 (2017).

49. Mitra, K., Wunder, C., Roysam, B., Lin, G. & Lippincott-Schwartz, J. A hyperfused mitochondrial state achieved at G<sub>1</sub>–S regulates cyclin E buildup and entry into S phase. Proceedings of the National Academy of Sciences 106, 11960–11965 (2009).

50. von Richter, O. et al. Cytochrome P450 3A4 and P-glycoprotein expression in human small intestinal enterocytes and hepatocytes: a comparative analysis in paired tissue specimens. Clinical pharmacology & therapeutics 75, 172–183 (2004).

51. Cheung, E.C. et al. Opposing effects of TIGAR-and RAC1-derived ROS on Wnt-driven proliferation in the mouse intestine. Genes & development 30, 52–63 (2016).

52. Friedman, E.S. et al. Microbes vs. chemistry in the origin of the anaerobic gut lumen. Proceedings of the National Academy of Sciences 115, 4170–4175 (2018).

53. Pfeiffer, T., Schuster, S. & Bonhoeffer, S. Cooperation and competition in the evolution of ATP-producing pathways. Science 292, 504–507 (2001).

54. Newsholme, E. & Carrie, A. Quantitative aspects of glucose and glutamine metabolism by intestinal cells. Gut 35, S13–S17 (1994).

55. Windmueller, H.G. & Spaeth, A.E. Identification of ketone bodies and glutamine as the major respiratory fuels in vivo for postabsorptive rat small intestine. Journal of Biological Chemistry 253, 69–76 (1978).

56. Castell, L., Bevan, S., Calder, P. & Newsholme, E. The role of glutamine in the immune system and in intestinal function in catabolic states. Amino acids 7, 231–243 (1994).

57. Arkblad, E.L. et al. A Caenorhabditis elegans mutant lacking functional nicotinamide nucleotide transhydrogenase displays increased sensitivity to oxidative stress. Free Radical Biology and Medicine 38, 1518–1525 (2005).

58. Sheeran, F.L., Rydström, J., Shakhparonov, M.I., Pestov, N.B. & Pepe, S. Diminished NADPH transhydrogenase activity and mitochondrial redox regulation in human failing myocardium. Biochimica et Biophysica Acta (BBA) - Bioenergetics 1797, 1138–1148 (2010).

59. Aw, T.Y. Intestinal glutathione: determinant of mucosal peroxide transport, metabolism, and oxidative susceptibility. Toxicology and Applied Pharmacology 204, 320–328 (2005).

60. Riddick, D.S. et al. NADPH–Cytochrome P450 Oxidoreductase: Roles in Physiology, Pharmacology, and Toxicology. Drug Metabolism and Disposition 41, 12–23 (2013).

61. Zhang, Q.Y., Wikoff, J., Dunbar, D. & Kaminsky, L. Characterization of rat small intestinal cytochrome P450 composition and inducibility. Drug Metabolism and Disposition 24, 322–328 (1996).

